# Nonself-recognition-based self-incompatibility can alternatively promote or prevent introgression

**DOI:** 10.1101/2020.09.29.318790

**Authors:** Alexander Harkness, Yaniv Brandvain

## Abstract

- Traditionally, we expect that self-incompatibility alleles (S-alleles), which prevent self-fertilization, should benefit from negative-frequency dependent selection and rise to high frequency when introduced to a new population through gene flow. However, the most taxonomically widespread form of self-incompatibility, the ribonuclease-based system ancestral to the core eudicots, functions through nonself-recognition, which drastically alters the process of S-allele diversification.
- We analyze a model of S-allele evolution in two populations connected by migration, focusing on comparisons among the fates of S-alleles originally unique to each population and those shared among populations.
- We find that both shared and unique S-alleles originating from the population with more unique S-alleles were usually fitter than S-alleles from the population with fewer. Resident S-alleles were often driven extinct and replaced by migrant S-alleles, though this outcome could be averted by pollen limitation or biased migration.
- Nonself-recognition-based self-incompatibility will usually either disfavor introgression of S-alleles or result in the whole-sale replacement of S-alleles from one population with those from another.

## 2 Introduction

In flowering plants, a self-incompatibility locus (S-locus) is a highly polymorphic region of the genome responsible for rejecting self pollen and is maintained by negative frequency-dependent selection, a form of balancing selection. A given S-locus allele (S-allele) encodes paired pollen and pistil phenotypes, and the pistil phenotype rejects pollen expressing the matching pollen phenotype, which necessarily includes self pollen. With self-incompatibility (SI), the fitness of an S-allele increases as it becomes rarer because rare alleles allow more mating opportunities. This process preserves polymorphism because alleles that drift to lower frequencies are pushed back to intermediate frequencies. It may also spur new polymorphism: new mutations are automatically rare, and their initial advantage under negative frequency-dependence favors their invasion (Wright, 1939; Charlesworth and Charlesworth, 1979). On first reflection, this advantage of rarity would seem to apply equally to novel mutations and rare migrant alleles. For one major form of self-incompatibility, self-recognition, the effects of this advantage of rarity have been modeled for both novel mutations (Uyenoyama et al., 2001) and rare migrants (Schierup et al., 1998; Muirhead, 2001; Schierup and Vekemans, 2008). But for nonself-recognition, the more taxonomically widespread form, only new mutations (Bod’ová et al., 2018) or, similarly, new products of gene conversion (Harkness et al., 2019) have been modeled. Here we ask if and when migration will lead to invasion of a new S-allele under nonself-recognition self-incompatibility. The answer to this work can influence both the-long term diversification of S-alleles and the role of S-allele divergence in preventing or promoting gene flow between diverged populations or species.

Invasion of a novel S-allele through rare advantage is simple if a single mutation can generate both a new pollen and pistil specificity (Charlesworth and Charlesworth, 1979). However, most self-incompatibility systems operate through linked but distinct pollen- and pistil-specificity loci, and a mutation at one component cannot affect the specificity encoded by the other. In such systems, S-alleles are more precisely called S-haplotypes, and while their maintenance by long term balancing selection is straightforward, the origin and diversification of novel two-locus S-haplotypes is complex. Uyenoyama et al. (2001) found in a deterministic two-mutation model that balancing selection in a single population could only increase S-haplotype diversity under stringent parameters and would usually lead either to loss of self-incompatibility entirely or to mere replacement of one S-haplotype with a novel one. They proposed that, though a local one-for-one replacement would not affect net local S-haplotype diversity of a subpopulation, all subpopulations collectively would retain the old haplotype while acquiring a new one. Subsequent gene flow could then spread the novel haplotype among subpopulations or reintroduce the old haplotype into the sub-population in which it was lost. Using extensive stochastic simulations of the same basic model, Gervais et al. (2011) found more relaxed conditions for local diversification but still invoked population structure to explain why natural populations harbor more haplotypes than were observed in their simulations.

But the foregoing are models of self-recognition based self-incompatibility, while the widespread S-locus ribonuclease (S-RNase) system ancestral to core eudicots functions through nonself-recognition (Kubo et al., 2010). Under self-recognition, each haplotype encodes a pollen specificity that is uniquely rejected by a pistil specificity on the same haplotype. This guarantees cross-compatibility of any two different S-haplotypes. The more complex nonself-recognition system can be envisioned as a lock-and-key system, in which each haplotype carries one pistil-specificity lock and many pollen-specificity keys (Harkness et al., 2019). Each lock can be unlocked by one key, and each haplotype carries a key ring that includes the key to every lock in the population but the haplotype’s own lock. Haploid pollen must carry keys to both the diploid seed parent’s locks in order to fertilize it. As such, nonself-recognition does not guarantee cross-compatibility with foreign specificities – as there is no reason to hold a key to a lock one never encounters. Therefore, just as Dobzhansky-Muller Incompatibilities (DMIs) (Dobzhansky, 1934; Muller, 1942) can accumulate between populations because the mutations with deleterious epistatic interactions are not normally exposed to one another, keys to locks present only in other populations are neutral and can easily be lost. Novel migrant haplotypes under nonself-recognition may therefore possess the advantage of rarity, just as under self-recognition, but they may also face disadvantages both as pollen parents – because they are incompatible with unique local haplotypes – and as seed parents – because they face a drastic reduction in compatible pollen.

We note that the dichotomy between self- and nonself-recognition is distinct from the better-known dichotomy between gametophytic (GSI) and sporophytic SI (SSI). In GSI, the phenotype of a pollen grain is completely determined by that pollen grain’s own haploid (gametophyte) genotype, while in SSI, the pollen phenotype is determined by that of the diploid (sporophyte) pollen parent. Different combinations of gametophytic or sporophytic and self- or nonself-recognition are possible: the S-locus receptor kinase system in Brassicaceae functions through sporophytic self-recognition, the programmed cell death system in poppy functions through gametophytic self-recognition, and the S-locus ribonuclease (S-RNase) system ancestral to the core euedicots functions through gametophytic nonself-recognition (Fujii et al., 2016). We are not aware of any examples of sporophytic nonself-recognition, but such a system might occur in any of the many taxa that have not been mechanistically characterized.

Recent models of the evolution of new S-haplotypes under non-self recognition, Bod’ová et al. (2018) and Harkness et al. (2019) found novel and surprising dynamics absent in self-recognition. “Complete” haplotypes, those that hold the keys to all other locks in the population and are thus compatible as pollen with all other haplotypes, have greater pollen success than “incomplete” haplotypes, which are missing some keys. An incomplete haplotype therefore cannot survive indefinitely in competition with more complete ones. Novel complete haplotypes may therefore eliminate existing incomplete haplotypes and actually reduce S-haplotype diversity unless subsequent mutation or gene conversion equalizes completeness among haplotypes. Stochastic simulations show that moderate numbers of haplotypes can be maintained through a continuous cycle of replacement in small or medium populations (Bod’ová et al., 2018), and deterministic approximations show that many haplotypes can be maintained in large populations or populations undergoing frequent gene conversion (Harkness et al., 2019).

Nonself-recognition thus presents unexpected challenges for the evolution of novel S-haplotypes, but what about the arrival of S-haplotypes through migration? Although both processes begin with an initially rare variant, the history of that rare variant is very different for each process. A novel S-haplotype generated by mutation or gene conversion is only one step removed from the other haplotypes in the same population: either it has added or lost a key on its key ring, or its lock has mutated such that it is unlocked by a different key. Whichever change occurs first, it is only the first step in a multi-step process requiring a new lock on one haplotype and a new key to that lock on every other haplotype. In contrast, a never-before-seen migrant haplotype has already completed this process: it has the keys to every lock in its own population, and all other haplotypes in its population have the key to its lock. Furthermore, there may be a consistent stream of migrant haplotypes every generation, whereas mutation or gene conversion presumably produce functionally novel haplotypes at a much lesser rate. These differences, the pre-existence of fully formed haplotypes and their continued introduction through migration, suggest that self-recognition might have very different implications for migration than for mutation or gene conversion.

We develop and compare population genetic models of gene flow under gametophytic self- and nonself-recognition to determine the effect of the widespread collaborative nonself-recognition on gene flow. We find that the general pattern under nonself-recognition is for gene flow to occur from the population with more unique haplotypes to the population with fewer, which may eliminate unique S-locus diversity in the recipient but may also facilitate the spread of novel alleles to multiple populations.

For the reader who would like more biological detail, we briefly present the molecular mechanism of S-RNase nonself-recognition SI here. Readers less interested in these details can skip to the **Description** section, which abstractly describes the haplotype cross-compatibility relationships necessary to understand the theoretical consequences of this system.

**The molecular mechanism of S-RNase nonself-recognition** is an area of ongoing research, and we merely present the leading hypothesis (reviewed by Williams et al., 2015), known as collaborative nonself-recognition (Kubo et al., 2010). Under this model, the S-locus consists of multiple tightly linked sites: a single highly polymorphic RNase gene and many biallelic (functional vs. non-functional/absent) S-locus F-box (SLF) genes around it (Kubo et al., 2010). The different SLF genes are diverged paralogs, but the allelic diversity at any one paralog is minimized by recurrent gene conversion (Kubo et al., 2015) despite the diversifying effect of long-term balancing selection among haplotypes. An SLF protein is expressed only in pollen and forms a part of the multi-protein SCF (Skp1-Cullin1-F-box protein) complex, a ubiquitin ligase (Lai et al., 2002; Qiao et al., 2004; Hua and Kao, 2006). Each diverged form of SLF protein targets this complex toward one or more allelic forms of RNase (Kubo et al., 2010). There are thus multiple versions of this complex, each targeted toward different RNases as determined by which SLF the complex possesses. The complex’s ultimate effect is to interfere with the RNases with which it interacts, though it is unclear whether this occurs through ubiquitin-labeling of the RNase for degradation by a proteasome or an alternative means (Hua and Kao, 2006).

The RNase is expressed in the style and transported inside the growing pollen tube regardless of whether the pollen is compatible or incompatible (Luu et al., 2000; Goldraij et al., 2006). If the pollen tube does not produce the SLF that targets this RNase, the RNase will slow or arrest pollen tube growth by digesting RNA in the pollen tube (Huang et al., 1994). Each maternal plant produces the two RNases in its diploid genotype (the metaphorical “locks”), so pollen is rejected by default unless it possesses the SLFs (the metaphorical “keys”) necessary to target both RNases. Self-incompatibility among copies of the same haplotype is achieved because each haplotype carries a nonfunctional or absent allele at the SLF paralog complementary to the haplotype’s own RNase, and cross-compatibility among different haplotypes is achieved because each haplotype carries a functional allele at the SLF paralogs complementary to all other RNases (Kubo et al., 2010).

## 3 Description

We modeled the evolution of the S-locus in two self-incompatible populations, the local population and the foreign population, of infinite size connected by pollen migration. In the simplest versions of this model, migration was unidirectional. From the perspective of the population receiving immigrants, that population is “local” while the population sending emigrants is “foreign.” For consistency, we retained this terminology throughout, but the local and foreign designations become arbitrary when migration is bidirectional. Each population harbored a number of S-haplotypes, some unique to that population and some shared between populations. We considered both self- and nonself-recognition models, but incompatibility was always gametophytic: the pollen phenotype was determined by its own haploid genotype, and the maternal phenotype was determined codominantly by its two haplotypes. Under GSI, the equilibrium condition in the absence of migration is equal frequency among S-haplotypes (Nagylaki, 1975; Boucher, 1993; Steiner and Gregorius, 1994), so we set this as the initial state for each population. Selection occurred only through pollen competition and maternal fecundity: there was no selection on viability. In pollen competition, all pollen compatible with a given maternal genotype competed equally to fertilize individuals of that maternal genotype. Pollen had zero success on maternal genotypes with which it was incompatible. We allowed for pollen limitation of maternal fecundity, in which maternal genotypes that accepted more pollen enjoyed greater seed success. Two considerations suggest that pollen limitation is an important process to model. First, since carriers of unique migrant haplotypes reject the majority of all pollen they receive (all resident haplotypes), they are likely to suffer severe pollen limitation in nature. Second, since pollen limitation is especially unfavorable to migrant haplotypes, we expect it to act as a barrier to gene flow that might partially counteract the advantage of rarity.

We use our lock and key metaphor (Harkness et al., 2019) to model non-self recognition. That is, the style is a door locked by two codominant RNase “locks,” and a pollen grain must carry the SLF “keys” to both of a seed parent’s locks in order to unlock the door and proceed to fertilization. For simplicity, we imagine that each key unlocks one lock, but in reality some SLFs are complementary to two or more RNases (e.g. Sun and Kao, 2013). Pollen is always incompatible with the plant that produced it because either possible key ring it could possess is missing one of the keys to one of the plant’s own two locks.

For self-recognition, we use the simplest model of a two-gene S-locus we could conceive, though we could not invent an adequate metaphor for this system. The S-locus under self-recognition consists of one polymorphic pistil-expressed gene tightly linked to one polymorphic pollen-expressed gene. Each pollen allele corresponds to one pistil allele, and corresponding alleles always exist on the same haplotype. By default, all pollen is accepted, but pollen carrying the pistil allele corresponding to either of the seed parent’s codominantly expressed pistil alleles is rejected. Since each haplotype carries a pollen allele that would be rejected by its own pistil allele, each haplotype is self-incompatible. This is similar to the programmed cell death mechanism in poppy, in which recognition between the pollen and pistil components triggers self-destruction in the pollen (Franklin-Tong et al., 1993), but differs from many better-known self-recognition based systems (Hiscock, 2002). We chose this system to isolate the effects of self vs nonself based recognition away from complications of dominance hierarchies among alleles in sporophytic SI systems.

Self- and nonself-recognition lead to different behavior in inter-population crosses. A self-recognition pistil allele only rejects one pollen allele: its complement. Non-matching pollen will always be accepted, regardless of its population of origin. In contrast, with nonself-recognition, a pistil locks out all pollen without a key. In a single population, this difference is immaterial: selection for maximal pollen success will ensure every S-haplotype is “complete” (Bod’ová et al., 2018) with respect to its own population, meaning that it carries keys to all locks in that population except its own lock. However, there would be little selection for a key to a lock that is rarely encountered and no selection for a key to a lock that is never encountered. Given enough time and isolation, populations will come to diverge in their sets of locks (either through novel mutations or differential loss), and each population will only maintain keys to locks present within that population. This leads to very different outcomes for haplotypes that are shared among populations and haplotypes unique to a population. Pollen from one population will always be compatible with any other haplotype from the same population, whether shared or unique. But pollen from one population will only be compatible with a haplotype from another population if that haplotype is shared with the pollen’s population of origin. We only explicitly model this ideal high-isolation case, but consider other plausible biological in the **Discussion**.

We classify all S-haplotypes based on their uniqueness and population of origin. This is sufficient for our model because the compatibility of pollen with locally unique S-haplotypes is determined by the pollen’s population of origin under nonself-recognition. S-haplotypes are either unique to the local population (*UL*), unique to the foreign population (*UF*), shared and originating from the local population (*SL*), or shared and originating from the foreign population (*SF*). Each class may contain one or more S-haplotypes, and we assume all haplotypes within a class always occur in equal proportions. We justify this assumption on the logic that, so long as it were true initially, it would remain true because haplotypes within the same class would have equal pollen and ovule success.

Shared local and shared foreign haplotypes come in pairs, e.g., a local *S*_1_ and a foreign *S*_1_. Under nonself-recognition, the members of a shared pair differ in their key rings: each has only the keys to locks originating from its own population. Under self-recognition, the members of a shared pair are functionally identical, though in nature they could be characterized by different neutral mutations. We use the same classification for self- and nonself-recognition for ease of comparison, but note that population of origin only affects phenotype under nonself-recognition, in which it determines an S-haplotype’s key ring. Functionally, there is only a single difference between our self- and nonself-recognition models: S-haplotypes originating from one population are incompatible as pollen with all haplotypes unique to another population under nonself-recognition but compatible under self-recognition (Fig. 1). This characterization allows us to compare the expected amount and pace of gene flow at the S-haplotype across comparable parameters for self- and non-self based SI.

**Figure 1:**
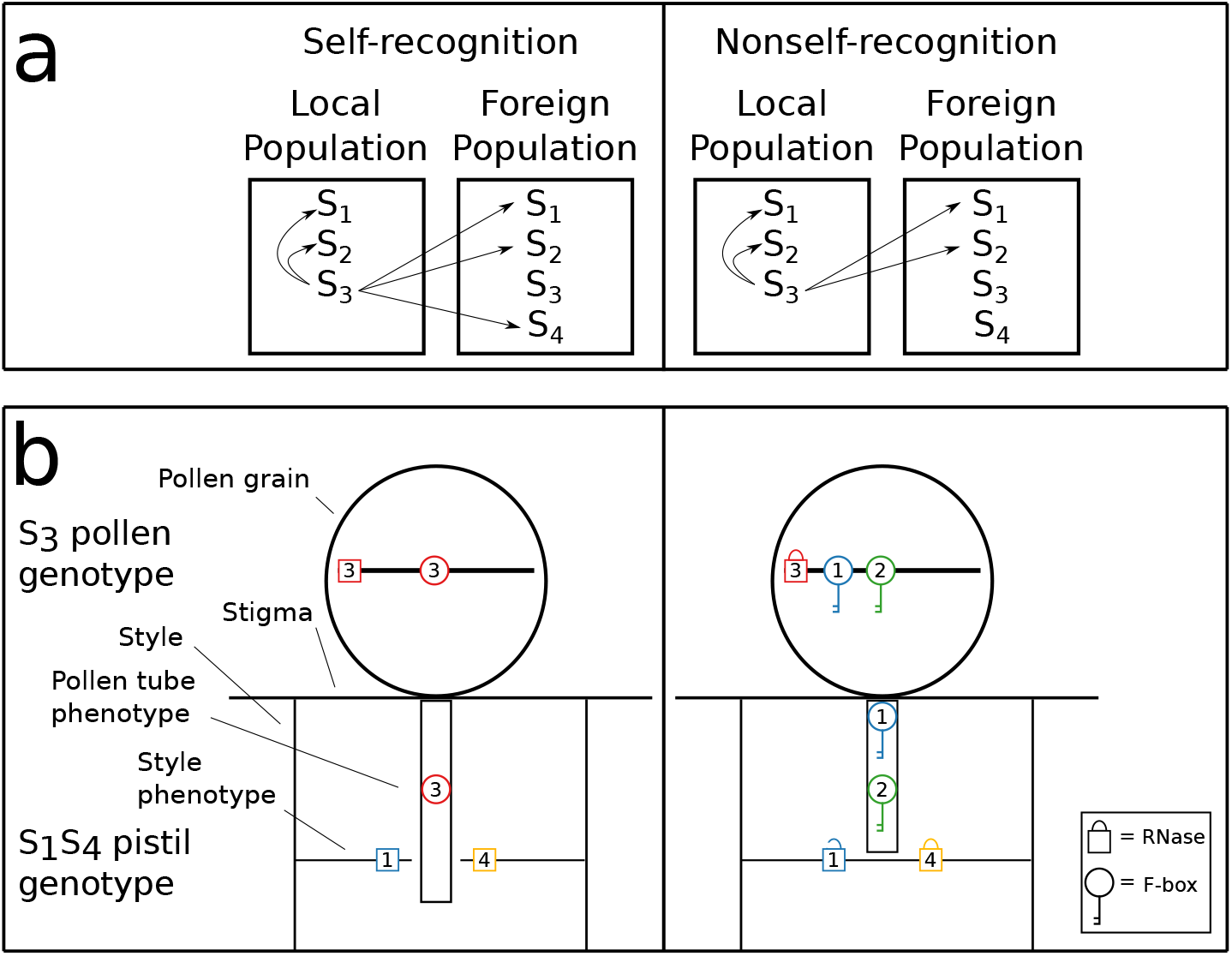
Pollen compatibility under self- and nonself-recognition. The focal *S*_3_ haplotype is shared and originates from the local population. *a* shows the consequences of self- and nonself-recognition for compatibility with other haplotypes. Under self-recognition, *S*_3_ pollen is only incompatible with *S*_3_-carrying plants. Under nonself-recognition, *S*_3_ pollen is still incompatible with *S*_3_-carrying plants, but it is also incompatible with carriers for haplotypes unique to another population (*S*_4_). *b* details the mechanistic basis of rejection for a local *S*_3_ pollen grain on a foreign *S*_1_*S*_4_ plant for self- and nonself-recognition. Under self-recognition, neither *S*_1_ nor *S*_4_ matches *S*_3_, so the pollen is not rejected. Under nonself-recognition, the local *S*_3_ haplotype’s key ring contains keys to all locks present in the local population (allowing it to unlock *S*_1_) but not the keys to locks unique to the foreign population (*S*_4_), so it is rejected. We place both rejection mechanisms in the style to match the RNase-based system, but rejection could occur at the stigma (as it does in some taxa) without affecting the model.

We denote the frequency of each class in each population by *p^L^* and *p^F^*. E.g., in the local population, the frequency of all haplotypes initially unique to the local population (*UL*) is 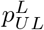. The frequency of a haplotype class is the sum of the frequencies of all haplotypes in that class. Since all haplotypes in a population begin at equal frequencies, the initial frequencies of the haplotype classes are determined by the number of haplotypes in each class. These numbers are denoted *n_L_* (number unique to the local population) *n_F_* (number unique to the foreign population), and *n_S_* (number shared). In the local population, initially,

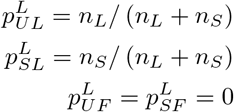

In the foreign population, initially,

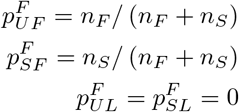

Gene flow occurs through pollen migration such that a proportion *m_FL_* of pollen in the local pollen pool is migrant pollen from the foreign population, and *m_LF_* of pollen in the foreign pollen pool of is migrant pollen from the local population. When migration is unidirectional from foreign to local, the foreign population remains at its initial equilibrium, so we only track the local population. When migration is bidirectional, we track both populations.

We model pollen limitation through a root function: a given genotype’s seed success is 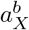, where *a_X_* is the frequency of all pollen compatible with genotype *X* in the available pollen pool and *b* is a shape parameter (with *b* = 0 corresponding to no pollen limitation). This allows seed success to saturate as compatible pollen received increases. Since the origin of a haplotype only affects is behavior in pollen, we drop the origin designation from the pollen accepted by each genotype *X*. E.g., *a_UFSF_* = *a_UFSL_*, so we instead simply write *a_FS_*.

We first describe analytical results for the case of unidirectional migration, and then describe numerical results for bidirectional migration. For this case, we focus entirely on frequencies within the local population and drop the superscript denoting population: e.g., the frequency in the local population of an allele initially unique to the foreign population 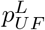 is simply labeled *p_UF_*. We use this case to lay out the three deterministic processes underlying haplotype frequency change.

First, selection on ovules occurs through pollen limitation. Each diploid genotype *X* contributes a number of successful ovules 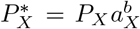. Second, assuming unidirectional migration, pollen migration generates the pollen pool. So

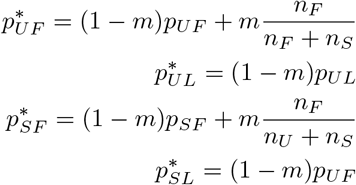

where a ^*^ denotes a frequency after migration.

Third, pollen selection occurs through differential pollen success, resulting in new genotype frequencies. Pollen fitness is a function of the proportion of stylar genotypes it can pollinate – which depends only on a pollen specificity’s frequency under self-recognition, and on the frequency of styles to which it holds the key under nonself recognition. These differing rules are reflected in the genotype frequency after mating under self- and nonself-recognition incompatibility (Appendix XX). We note that with nonself-recognition, no individual ever carries both a unique foreign and unique local 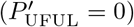 because neither holds the key to the other’s lock. We further note that these analytical results did not include pollen limitation, a biologically relevant process which we explore in our numeric iterations.

We derive analytical results for a special case. Focusing on nonself-recognition and with no pollen limitation, the marginal fitnesses of the shared haplotypes are

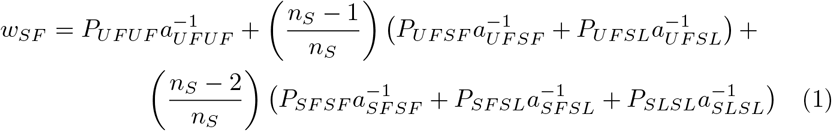

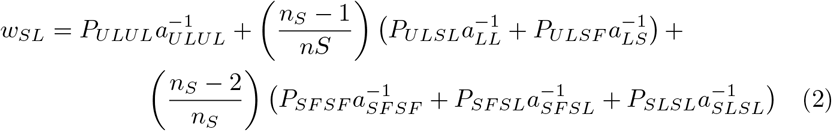

Taking the difference, the third term in each cancels out,

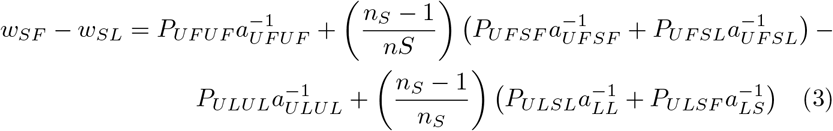

and all remaining terms depend on the frequency of carriers for the unique haplotypes. The sign of this difference is obvious when one population harbors no unique haplotypes. For example, when the local population has no unique S-haplotypes (*n_L_* = 0) and the foreign population has at least one unique S-haplotype (*n_F_* > 0), the fitness of shared haplotypes from the foreign population exceeds that of shared haplotypes from the local population. Since *w_SL_ < w_UF_* at all genotype frequencies, there is no non-zero equilibrium frequency of *p_SL_*. So long as *p_SF_* > 0 and *p_UF_* > 0, the *SL* haplotype will be eliminated at equilibrium. Similarly, when *n_L_* > 0 and *n_F_* = 0, *w_SL_ > w_SF_* at all genotype frequencies.

The biological interpretation of this result is straightforward. The only difference between shared haplotypes originating from different populations is their collection of pollen-function keys: all *L* haplotypes (shared or unique) are compatible with S-haplotypes initially unique to the local population (*UL*), while *SF* haplotypes are compatible with *UF* haplotypes. If *UF* haplotypes exist but *UL* haplotypes do not, *SF* haplotypes have all the pollen success of *SL* haplotypes plus additional pollen success on *UF* haplotypes. In this case, each *SF* haplotype is always fitter than its *SL* counterpart, and selection will drive all *SL* haplotypes extinct unless opposed by biased migration. That is, a population harboring no unique haplotypes will have its shared haplotypes replaced by shared haplotypes from the other population. No such process occurs under self-recognition.

This analytical result suggests an important role for the relative numbers of unique haplotypes in each population. To investigate the effect of these and other parameters in more general cases, we implemented the full deterministic model in R. Using this model, we tracked evolution of haplotype frequencies under scenarios varying the numbers of shared (*n_S_*) and unique haplotypes (*n_L_* and *n_F_*), the presence (*b* = 1*/*2) or absence of pollen limitation (*b* = 0), the duration of migration (continuous or a single pulse), and the form of self-incompatibility (self- or nonself-recognition). For this part of the investigation, we kept migration unidirectional from foreign to local (*m_LF_* = 0). Migration persisted throughout the iterations if continuous or ceased after a single generation if pulsed, and it occurred at the same modest rate (*m_FL_* = 0.01) whether continuous or pulsed.

After observing that nonself-recognition sometimes facilitated and sometimes prevented introgression of migrant haplotypes, we quantified the minimum value of *m_FL_* that resulted in invasion of *SF* for different values of *n_L_* and *n_F_* under nonself-recognition. We varied *m_FL_* from 0–0.1 in increments of 0.01 and varied *n_L_* and *n_F_* from 0–10. We considered values of *n_S_* = 5 or 20 and *b* = 0 or 1/2 but held *m_LF_* = 0 constant. Invasion was considered to have occurred if *p_UF_* + *p_SF_ > p_UL_* + *p_SL_* in the local population at generation 1000. Note that this definition of invasion does not imply positive selection on the *SF* haplotypes: a neutral haplotype would also invade under this unidirectional migration scheme.

Finally, we tracked haplotype frequencies for several bidirectional migration scenarios. We considered four scenarios in which migration and the number of shared haplotypes were held constant (*m_LF_* = *m_FL_* − 1, *n_S_* = 5), in which one (*n_L_* = 0 and *n_F_* = 1) or both populations possessed a unique allele (*n_L_* = *n_F_* = 1), and in which pollen limitation was either present (*b* = 1/2) or absent (*b* = 0). Additionally, we more closely investigated the haplotype freqeuncy trajectories in another four scenarios, which had resulted in coexistence of foreign and local haplotypes at equilibrium. In these scenarios, the numbers of shared and unique haplotypes and the strength of pollen limitation were constant (*n_S_* = 20, *n_L_* = *n_F_* = 1, *b* = 1*/*2), and migration was either low and symmetric (*m_LF_* = *m_FL_* = 0.02), high and symmetric (*m_LF_* = *m_FL_* = 0.09), asymmetric with more immigration to the local population (*m_LF_* = 0.05, *m_FL_* = 0.06), or asymmetric with more immigration to the foreign population (*m_LF_* = 0.05, *m_FL_* = 0.04).

## 4 Results

### Self-recognition-based SI

The consistent outcome of self-recognition was for rare pistil specificities to rise to higher frequencies regardless of their origin (Fig. 2). All unique haplotypes always coexisted at equilibrium for both ongoing unidirectional migration (Fig. 2A,B) and a one time migration pulse (Fig. 2C,D).

**Figure 2:**
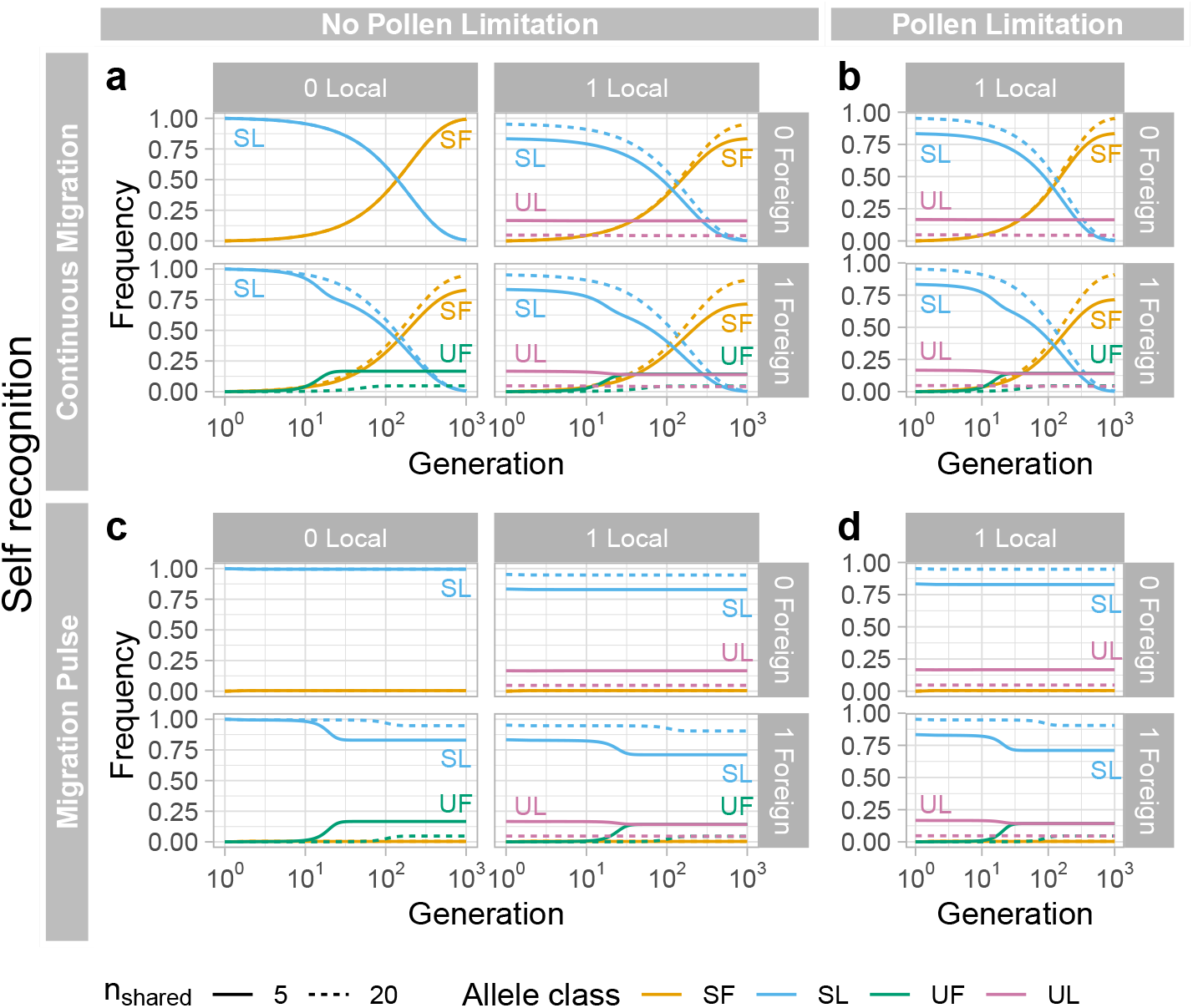
Evolutionary dynamics of shared foreign (orange), shared local (blue), unique foreign (green), and unique local (pink), S-haplotypes with self-recognition based SI. Results for a steady (unidirectional) migration rate of 0.01/generation (*2A* and *2B*). Results for a one time migration pulse 0.01 (*2C* and *2D*). We show results without (*2A* and *2C*) and with (*2B* and *2D*) pollen limitation. Facets atop and beside the plots show the number of unique local and foreign haplotypes, respectively. Dashed and full lines show the cases of twenty and five shared haplotypes, respectively. Generations increase on a *log*_10_ scale on the x-axis.

Under self-recognition, pollen limitation did not severely impact the spread of foreign S-haplotypes (c.f. the right hand columns of Fig. 2A and Fig. 2C, to Fig. 2B and Fig. 2D) because styles of local and foreign shared and unique haplotypes can accept all nonself pollen. Rather, the equilibrium frequency of *SL* and *SF* haplotypes was determined entirely by migration pressure. Under continuous unidirectional migration, shared haplotypes originating in the donor population globally displaced shared haplotypes originating in the recipient population (Fig. 2). But under a single pulse of unidirectional migration, when only selection was ongoing, the initially low-frequency unique haplotypes migrating from the foreign population (*UF*) rose to match the frequency of the haplotypes unique to the local population (*UL*), while shared haplotypes (*SF*) remained at their initial low frequency (Fig. 2).

We note that the equilibrium frequency of all shared haplotypes relative to all unique haplotypes increased as the number of shared haplotypes increased (compare dashed and full lines in Fig. 2), as would be expected from increasing the number of haplotypes in a category and counting them collectively. A similar result also occurs with nonself-recognition, as seen in Fig. 3. Because results from the case of self recognition are straightforward and confirmed current understanding, we spend the remainder of our efforts on the results from nonself-recognition SI.

**Figure 3:**
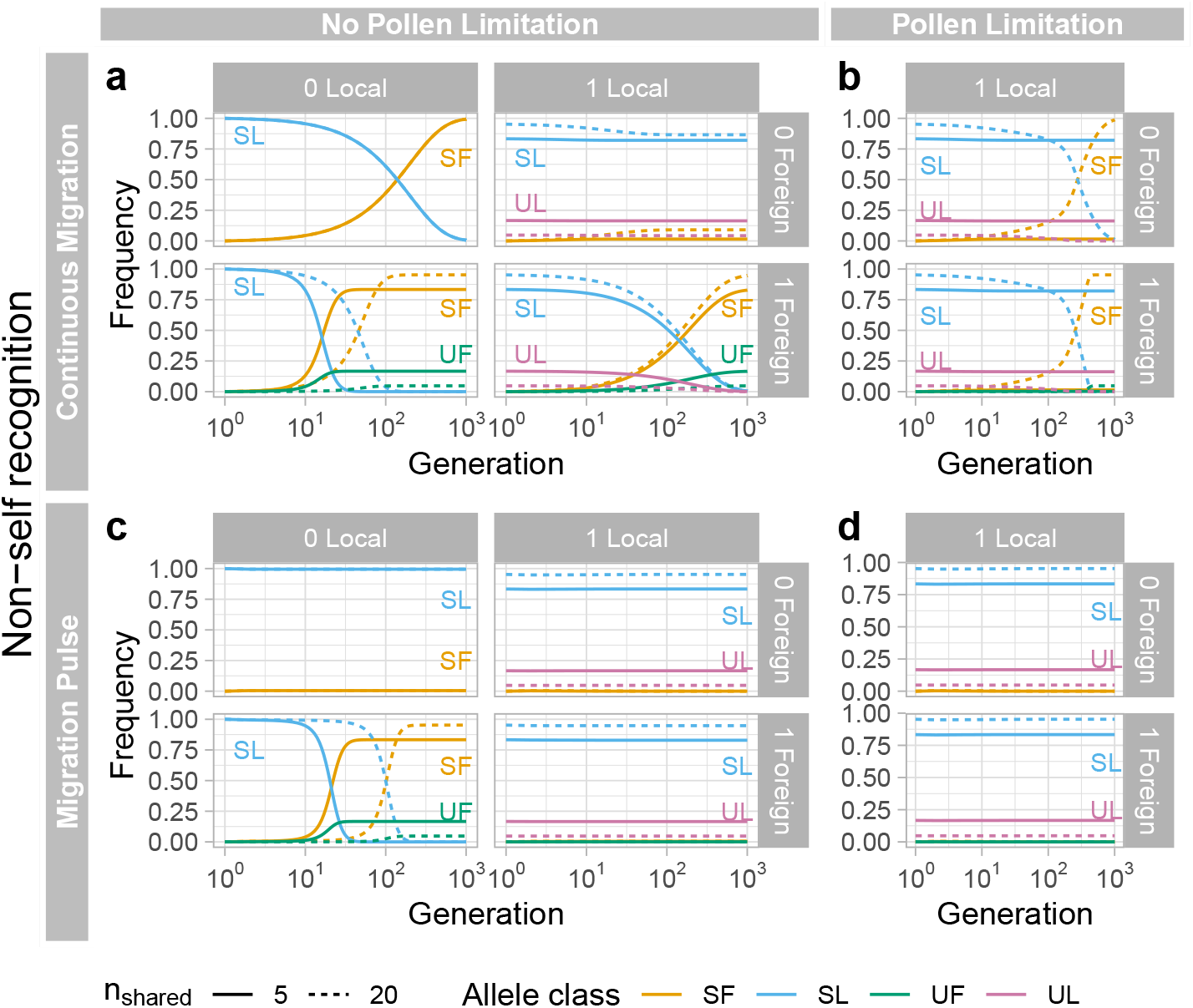
Evolutionary dynamics of shared foreign (orange), shared local (blue), unique foreign (green), and unique local (pink), S-haplotypes with nonself-recognition based SI. Results for a steady (unidirectional) migration rate of 0.01/generation (*2A* and *2B*). Results for a one time migration pulse 0.01 (*2C* and *2D*). We show results without (*2A* and *2C*) and with (*2B* and *2D*) pollen limitation. Facets atop and beside the plots show the number of unique local and foreign haplotypes, respectively. Dashed and full lines show the cases of twenty and five shared haplotypes, respectively. Generations increase on a *log*_10_ scale on the x-axis. Figure layout is identical to that in Figure 2.

### Nonself-recognition-based SI

Under nonself-recognition, the results are sensitive to parameter values, and many outcomes were possible.

Without pollen limitation and with ongoing unidirectional gene flow, foreign unique S-haplotypes can often invade. For example, with one foreign and no local unique S-haplotypes, the unique foreign S-haplotype rapidly establishes and reaches its equilibrium frequency (Bottom row, first column of Fig. 3A). This process is somewhat slower when the local population also has a unique S-haplotype (Bottom row, second column of Fig. 3A). Unlike the case of self-recognition, with nonself-recognition, the foreign unique S-haplotype replaces the local unique S-haplotype (when green lines in e.g. the bottom panel of Fig. 3A are above zero, pink lines head towards zero). Additionally, because pollen of both local shared and local unique S-haplotypes is incompatible with styles with unique foreign locks, its fitness decreased as the unique foreign S-haplotype rose in frequency, consistent with our analytical predictions (Eq. 3). In contrast, the shared foreign S-haplotype enjoyed rapidly increasing pollen fitness as the unique foreign haplotype rose in frequency (compare the slower rise in frequency of shared foreign haplotypes in the bottom row of Fig. 3A to that in Fig. 2A). As a result, foreign shared haplotypes replaced local shared haplotypes, just as foreign unique haplotypes replaced local unique haplotypes.

In the absence of pollen limitation, increasing the number of shared haplotypes from 5 to 20 did not qualitatively affect the outcome of migration but modified the equilibrium frequency of shared and unique haplotypes (compare dashed and full lines Fig. 3A), as observed in the case of self-recognition, above. By contrast, the number of shared S-haplotypes mediated the spread of foreign S-haplotypes when there was pollen limitation. For example, in Fig. 3B, foreign haplotypes cannot invade and replace local haplotypes when there are five shared haplotypes but can when there are twenty. This difference reflects the extent of pollen limitation faced by unique foreign haplotypes – within twenty shared haplotypes, a plant with a unique foreign S-haplotype can be fertilized by 95% (20 of 21) of local pollen haplotypes, a large increase from the 83% (5 of 6) in the case of five shared S-haplotypes.

Despite long-term balancing selection favoring the proliferation of S-haplotypes, the initial disadvantages of rejecting most local pollen as a maternal plant and being rejected as pollen by unique local haplotypes can disfavor rare foreign haplotypes. As such, although foreign S-haplotypes displace local ones with on-going migration (Figures 3A,B), a single small pulse of migration does not result in the invasion of foreign haplotypes unless there are no unique local haplotypes (Figures 3C,D) under nonself-recognition.

To quantify the effects of shared and unique haplotypes, as mediated by the extent of gene flow, we calculated the minimum value of *m_FL_* for which *p_UF_* + *p_SF_ > p_UL_* + *p_SL_* at generation 1000 (Fig. 4). Without pollen limitation, invasion was possible for a broad range of *n_L_* and *n_F_* values, though it could occur even at low migration rates when *n_F_ ≥ n_L_*. With pollen limitation and *n_S_* = 5 shared haplotypes, invasion was only possible for *m_FL_* ≤ 0.1 when *n_F_ ≥ n_L_*. However, the exact effect of pollen limitation depended on *n_S_*. At *n_S_* = 5, pollen limitation consistently raised the threshold migration for invasion. At *n_S_* = 20, pollen limitation usually raised the threshold when *n_L_ < n_F_* but lowered it when *n_L_ > n_F_*. When pollen is limiting and there is some migration, the seed success of unique haplotypes is reduced. When there are many haplotypes, the pollen advantage of rarity is smaller. These disadvantages combined may be sufficient to eliminate rare unique haplotypes, thereby eliminating a barrier to the invasion of foreign haplotypes whether shared or unique. We demonstrate such a process for a case of bidirectional migration below.

**Figure 4:**
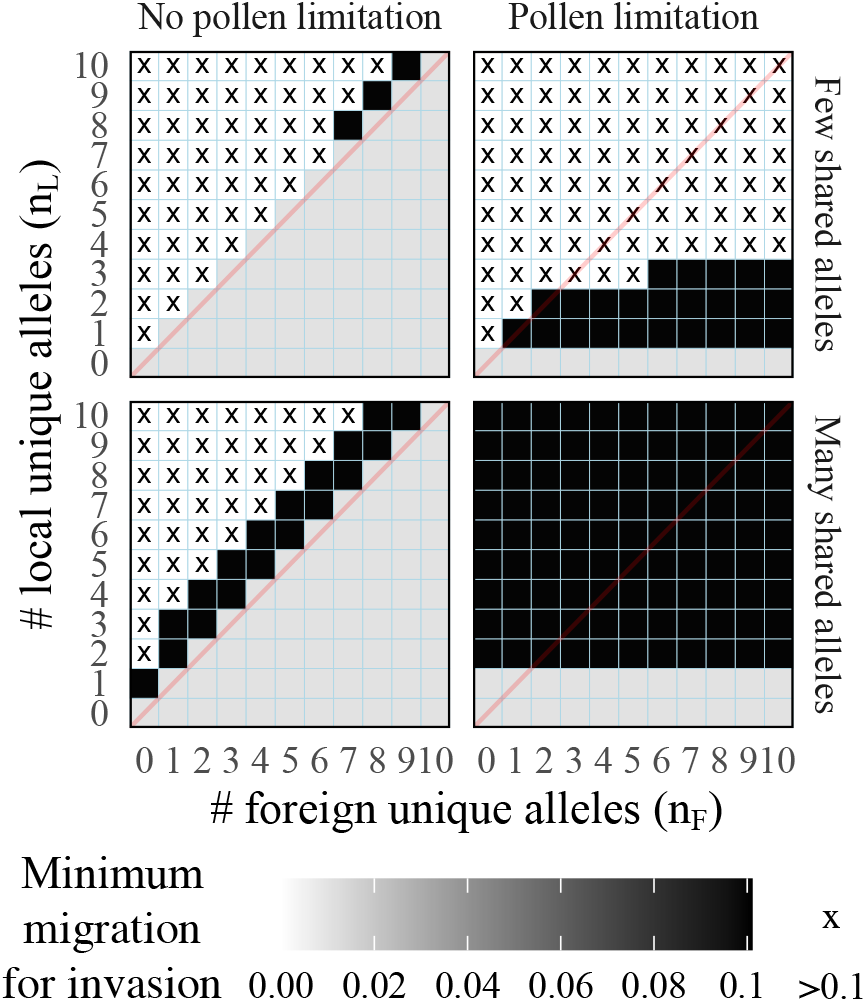
Effect of unique haplotypes on invasion threshold. The minimum rate of unidirectional migration (*m_FL_*) needed for the frequency of all foreign haplotypes to exceed the frequency of all local haplotypes (*p_UF_* + *p_SF_ > p_UL_* + *p_SL_*) in the local population at generation 1000. Invasion could occur at low migration rates when the number of foreign unique haplotypes (*n_F_*) exceeded the number of local unique haplotypes (*n_L_*) but required high migration rates when *n_L_ > n_F_*. Many shared haplotypes (*n_S_* = 20) lowered the invasion threshold compared to few shared haplotypes (*n_S_* = 5). Pollen limitation raised the invasion threshold when *n_S_* = 5, but when *n_S_* = 20, the effect of pollen limitation was to erase partly the effect of *n_F_*. Seed set for a genotype was a square root function of the proportion of pollen compatible with that genotype. The *y* = *x* line is shown in faint red.

When migration was equal in both directions, the equilibrium state depended mainly on the numbers of unique haplotypes. If only one population possessed a unique haplotype, the shared haplotype from the same population globally replaced the shared haplotype from the other population alongside the invading unique haplotype (Fig. 5). If both populations possessed a unique haplotype, both populations converged on the same equilibriumin the absence of pollen limitation or maintained their initial states in the presence of pollen limitation (Fig. 5).

**Figure 5:**
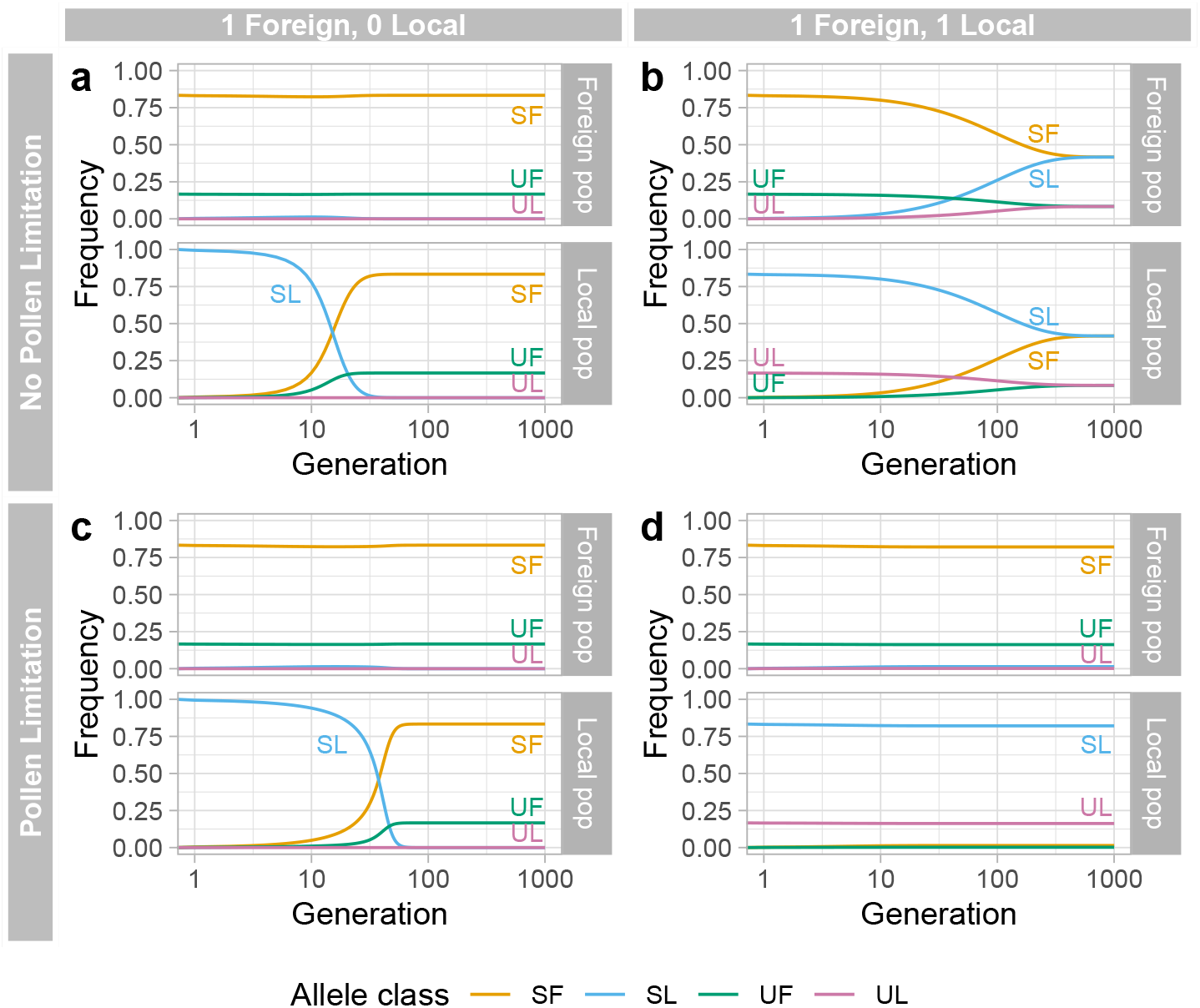
Bidirectional migration. Migration rates are *m_FL_* = *m_LF_* = 0.01. There were *n_S_* = 5 shared haplotypes. All local haplotypes were lost when only the foreign population possessed a unique haplotypes, regardless of pollen limitation (A, C). When both population possessed one unique haplotype, local and foreign haplotypes coexisted either by converging toward the same intermediate equilibrium frequencies in both populations in the absence of pollen limitation (B) or by being preserved at high frequencies in their original populations in the presence of pollen limitation (D).

Under broader ranges of migration rates, including unequal ones, four results were possible: elimination of local haplotypes, elimination of foreign haplotypes, coexistence of foreign and local haplotypes within each population, or maintenance of each unique haplotype only in its population of origin (Fig. 6). The population sending more emigrant pollen tended to preserve its haplotypes. Without pollen limitation, coexistence occurred easily, but the exact equilibria of foreign and local haplotypes depended on migration rates. With pollen limitation, coexistence only occurred in a narrower band of intermediate migration rates. In most cases, both populations converged on the same equilibrium as long as migration was nonzero in either direction. But in the presence of both pollen limitation and few shared haplotypes, each population could maintain its own unique haplotype at high frequency while preventing much introgression of the other population’s unique haplotype. Sufficiently biased migration could overcome this maintenance of distinct haplotypes and allow one unique haplotype to replace the other globally.

**Figure 6:**
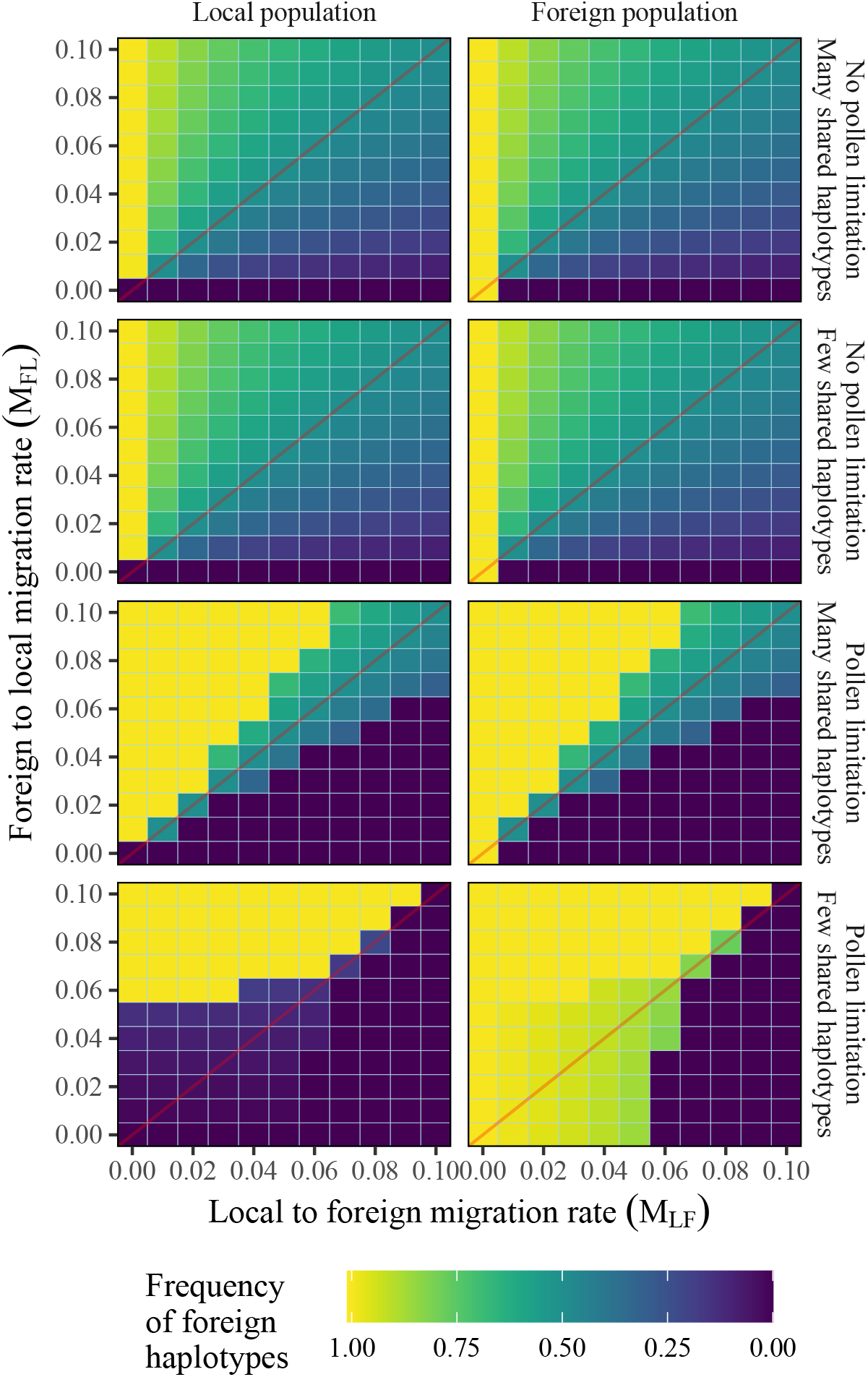
Equilibrium frequency of foreign haplotypes in each population with bidirectional migration. Self-incompatibility functions through nonself-recognition in all panels. Each population harbors one unique haplotype (*n_L_* = *n_F_* = 1), and many (*n_S_* = 20) or few (*n_S_* = 5) shared haplotypes. In the absence of pollen limitation, foreign and local haplotypes usually coexist unless migration rates are extremely biased. With pollen limitation, moderately biased migration rates result in the loss of foreign or local haplotypes, and co-existence is only possible for a narrower band of more nearly equal migration rates.

In some cases, pollen limitation could drive unique haplotypes from both populations extinct (Fig. 7). Since unique haplotypes could not accept immigrant pollen, they suffered reduced seed success compared to shared haplotypes. Once the unique haplotypes were gone, the distinction between shared haplotypes from different populations became irrelevant to selection: each kind carried a key to a different extinct lock. The shared haplotypes could then coexist in the absence of unique haplotypes.

**Figure 7:**
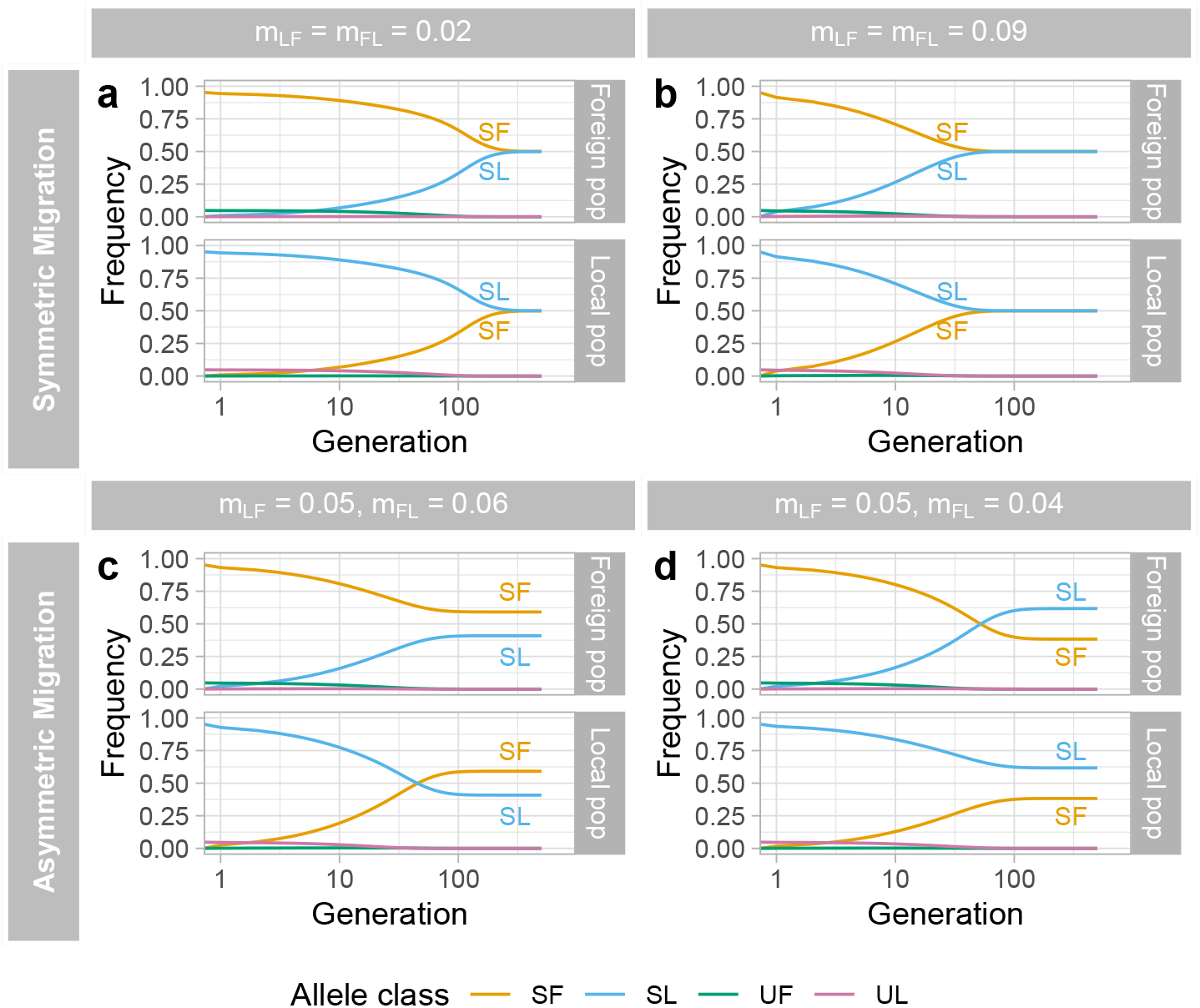
Loss of diversity. At some intermediate migration rates, shared haplotypes (*SL* and *SF*) can coexist while all unique haplotypes (*UF* and *UL*) are lost. This outcome can occur for both symmetric (A, B) and asymmetric (C, D) migration rates, with asymmetric rates altering the equilibrium frequencies of the shared alleles. There were *nS* = 20 shared haplotypes, *n_L_* = 1 unique local haplotype, and *n_F_* = 1 unique foreign haplotype, and seed success was a square root function of compatible pollen.

## 5 Discussion

Existing theory predicts that rare advantage at an S-locus will elevate gene flow by favoring rare immigrant S-haplotypes (e.g., Pickup et al., 2019). We confirm this prediction for self-recognition systems, but find that nonself-recognition can either enhance or reduce gene flow depending on the S-haplotypes present in each population. The crucial parameter is the number of S-haplotypes unique to each population because pollen originating from one population is incompatible with S-haplotypes unique to the other. Unique S-haplotypes can act as a barrier to incoming gene flow by rejecting foreign pollen. At the same time, they can enhance outgoing gene flow because, once they enter the other population, they reject resident pollen and reduce its fitness relative to immigrant pollen. We find that introgression of S-haplotypes from the population with more S-haplotypes to the population with fewer is usually elevated, while introgression in the reverse direction is usually reduced. If this haplotype-based bias points in the same direction as the bias in migration, or if migration is unbiased, introgression is elevated. But the haplotype-based bias instead impedes introgression if it is opposite to the direction of migration bias. We should therefore expect very different patterns of gene flow at the S-locus in taxa with nonself-recognition, the ancestral state in core eudicots (Igic and Kohn, 2001), and taxa that have evolved some form of self-recognition, as in Papaveraceae and Brassicaceae.

Importantly, biased introgression under at the S-locus under non-self recognition applied to shared haplotypes as well as unique haplotypes. As a unique migrant haplotype invades, the advantage of compatibility with that haplotype increases. Thus, a unique haplotype can invade by the advantage of rarity while the shared haplotypes from the same population ride its coattails. A common pattern we observed was that the population with more unique haplotypes or higher emigration rates essentially swamped the other population, replacing both unique and shared haplotypes.

The role of introgressed locks in facilitating the invasion of keys originating from the same population is analogous to the surprising effect of introgressed female preferences on male traits in sexual selection predicted by Servedio and Bürger (2014). Sexual selection has classically been viewed as a barrier to introgression because, if two populations have divergent female preferences, migrant males with out-of-place traits will suffer reduced reproductive fitness. However, female preferences may themselves introgress if they not under direct selection. As a preference introgresses, the reproductive fitness of the corresponding male trait increases, allowing the trait to increase in frequency. This facilitated introgression can counteract the fitness advantage of so-called “magic traits” that are both adaptive in the local environment and initially preferred by local females. Pistil-expressed self-incompatibility locks are analogous to strong female preferences for a given pollen-expressed key, analogous to a male trait. The difference is that, instead of one male trait with different values in each population, pollen keys are a collection of many binary traits. Thus, pollen with more keys may be preferable to many seed parents in both populations, resulting in a directional asymmetry absent from pure Fisherian sexual selection.

Pollen limitation and the number of shared haplotypes also affected introgression of foreign S-haplotypes. Pollen limitation typically reduced introgression and could result in the maintenance of two effectively isolated populations with their own sets of haplotypes. In contrast, introgression increased as the number of shared haplotypes increased. Shared haplotypes, unlike unique haplotypes, could accept immigrant pollen. When there were many shared haplotypes, immigrant pollen was rarely rejected and had high fitness.

Surprisingly, when migration was bidirectional and roughly symmetric, pollen limitation could instead eliminate unique haplotypes in both populations, which reduced the overall number of haplotypes but allowed shared haplotypes to introgress freely. Unique haplotypes were lost because they rejected migrant pollen and thus suffered greater pollen limitation than shared haplotypes, which did not. This is a consequence of the model’s square root function of pollen limitation, in which seed success monotonically increases with compatible pollen and never fully flattens. Therefore, accepting more pollen always increases seed success somewhat, and unique haplotypes are disadvantaged as seed parents. If seed success instead plateaus, unique haplotypes should still be disadvantaged unless both shared and unique haplotypes accept enough pollen to reach the plateau.

From a genomic perspective, each invasion of a unique migrant haplotype under nonself-recognition should reduce among-population neutral divergence for all haplotypes at the S-locus. In contrast, invasion of a migrant haplotype under self-recognition should only reduce divergence among copies of a single haplotype. We therefore predict greater among-population neutral divergence at the S-locus in species with self-recognition than in those with nonself-recognition. Surprisingly, a greater rate of S-haplotype diversification might therefore lead to reduced divergence between populations at the S-locus. Data on S-allele overlap and divergence across populations is available, at least for self-recognition. In *Arabidopsis halleri* and *A. lyrata*, which possess the self-recognition-based S-locus receptor kinase (SRK) incompatibility system, Castric et al. (2008) found 18 pairs of SRK alleles diverging by less than or equal to 12 substitutions. The remaining 12 alleles in *A. halleri* and 20 alleles in *A. lyrata* had no such counterparts in the other population. The low divergence within each pair could be explained by elevated gene flow of these alleles but not by the mere reduced effective population size of a single S-allele. This pattern is consistent with our expectation for self-recognition that migrant S-alleles will rise to high frequency without greatly reducing diversity in the S-locus as a whole. However, we also predict that any pairs of functionally identical S-alleles that were initially shared by the two species would have persisted in each population. It is possible that these originally shared alleles have since been lost in one or both populations or that they have since accumulated enough neutral or functional divergence that they are no longer recognizably shared. We also note that the SRK system is sporophytic rather than the gametophytic self-recognition system we modeled.

More comprehensive theory already exists for the interaction between haplotype number and migration in the case of self-recognition. In a model combining migration and balancing selection (either symmetrical overdominance or SI), Muirhead (2001) predicted for a given migration rate both the expected total number of alleles and the distribution of the proportion of alleles shared between two populations, three populations, etc. For the total number, she found a non-monotonic relationship in which allele number is minimized for intermediate migration rates. This non-monotonic relationship was previously observed in simulations by Schierup (1998). For the proportions of shared alleles, she found that increasing the migration rate skewed the distribution towards alleles shared among many populations. In the SI version of this model, all unlike S-haplotypes were assumed to be cross-compatible. This is equivalent either to nonself-recognition in which all haplotypes are complete or to self-recognition. However, it is not equivalent to nonself-recognition in which some haplotypes are more complete than others.

Theory on both self- and nonself-recognition has revealed hurdles to S-haplotype diversification and suggested that gene flow can help populations overcome these hurdles (Uyenoyama et al., 2001; Gervais et al., 2011; Harkness et al., 2019). Our results constrain how S-haplotype diversification could occur in a subdivided population or metapopulation. Uyenoyama et al. (2001) found that local turnover, replacement of old haplotypes without increasing the total number, was possible under a much broader set of parameter values than local diversification. They therefore hypothesized that diversification occurs through local turnover followed by introgression of the new haplotype into the metapopulation and reintroduction of the lost haplotype from the metapopulation. We predict under this process that novel S-haplotypes from the same population can easily spread simultaneously, but novel haplotypes from different populations may interfere and eliminate each other. Reintroduction of a haplotype lost to turnover would require the reintroduced haplotype to become compatible with the novel haplotype. Until cross-compatibility is restored, the novel haplotype may continue to replace the locally lost haplotype in every subpopulation. Gene conversion could restore cross-compatibility (Kubo et al., 2015; Fujii et al., 2016), potentially halting the loss of a haplotype (Bod’ová et al., 2018; Harkness et al., 2019).

Although we isolated effects of migration, in nature it should operate simultaneously with mutation and gene conversion. There may be a separation of time scales such that migration resolves itself before new mutations or gene conversions can occur, but certain mutations or gene conversions might push the system toward a new equilibrium. Some of the most common events might be those that induce self-compatibility: either the lock suffers a loss-of-function mutation or a haplotype acquires the key to its own lock by gene conversion. Mutant haplotypes with a nonfunctional lock would reject no pollen and would reduce pollen limitation, and these self-compatible mutants might introgress more easily than self-incompatible haplotypes. In contrast, a haplotype with the key to its own lock would only affect the pollen phenotype: self pollen would be compatible, but the maternal plant would still reject the same subset of nonself pollen. While acquiring a key (which offers new siring opportunities) would typically increase fitness more than losing a lock, the substantial pollen limitation suffered by migrant haplotypes might reverse this inequality: losing a lock would completely eliminate pollen limitation by accepting all pollen, while gaining a key would only mitigate pollen limitation by accepting self pollen. If accepting self pollen is insufficient to achieve full seed set, losing a lock should provide greater benefit to seed set than gaining a key.

We assumed that all haplotypes were incompatible with haplotypes unique to other populations. But if the difference in S-haplotypes between populations was caused by recent differential loss, it is likely that each population would temporarily retain cross-compatibility with locally lost haplotypes. That is, the loss of a pistil-function lock does not imply immediate loss of the corresponding pollen-function key from all other haplotypes. However, mutations to this now-neutral key would eventually render it non-functional. Another possibility is that, if two pistil specificities correspond to the same dual-function pollen key, cross-compatibility with a foreign pistil lock might have been maintained as a byproduct of selection for compatibility with a local pistil lock. Furthermore, even if foreign haplotypes are initially rejected by local unique haplotypes, the foreign haplotypes could acquire the missing local key through gene conversion with local shared haplotypes. If a haplotype is being driven extinct, it may be rescued by acquiring a missing key and increasing its pollen fitness (Bod’ová et al., 2018; Harkness et al., 2019).

We have only modeled the S-locus, but any S-haplotype in nature comes as part of a whole parental genome. That genome might not be as well adapted to the local environment as resident genomes. In this case, linkage disequilibrium between the S-locus and loci involved in environmental adaptation might slow the introgression of migrant S-haplotypes. Recombination can break down this linkage disequilibrium in the long run, but there may be few opportunities to recombine off a locally maladapted background if hybrids are rare. Gene flow at the S-locus, like other loci, should be limited by the density of locally adapted loci around it.

The discovery of collaborative nonself-recognition has unexpected theoretical implications for the behavior of the S-locus. Recent theory on S-haplotype diversification reveals complicated dynamics of collapse and rescue (Bod’ová et al., 2018; Harkness et al., 2019). Similarly, we show that gene flow is more complex under nonself-than under self-recognition. The S-locus, an ancient and widespread feature controlling the breeding system of many flowering plants, seems only to get stranger the more it is investigated.

## 6 Acknowledgments

This work was supported by **NSF DEB #1754246 to Y. B.**

A. H. was funded through the University of Minnesota Doctoral Dissertation Fellowship. We would like to thank Dr. Shelley Sianta and Dr. Emma Goldberg, whose especially thorough and insightful feedback greatly improved the rigor, clarity, and focus of this manuscript.

## 7 Author Contribution

A. H. conceived the project, designed and analyzed the model, wrote each major draft, and produced initial exploratory figures.

Y. B. revised and edited drafts, provided feedback as the model was developed, and created all final data visualizations.

## 8 Appendix

## 8.1 Nonself-recognition

After pollen migration, the frequency of each haplotype in the pollen pool is changed from *p* to *p*^*^. For shared haplotypes, population of origin only affects pollen behavior. Therefores, there are effectively only three kinds of haplotypes from the perspective of maternal plant phenotype: local unique haplotypes, foreign unique haplotypes, and all shared haplotypes collectively. We drop the population of origin when denoting the pollen accepted by a genotype: e.g., both *UF SF* and *UF SL* accept the same proportion *a_FS_* of pollen rather than distinct values of *a_UFSF_* and *a_UFSL_*. Assuming no pollen limitation, mating with nonself-recognition produces the genotype frequencies:

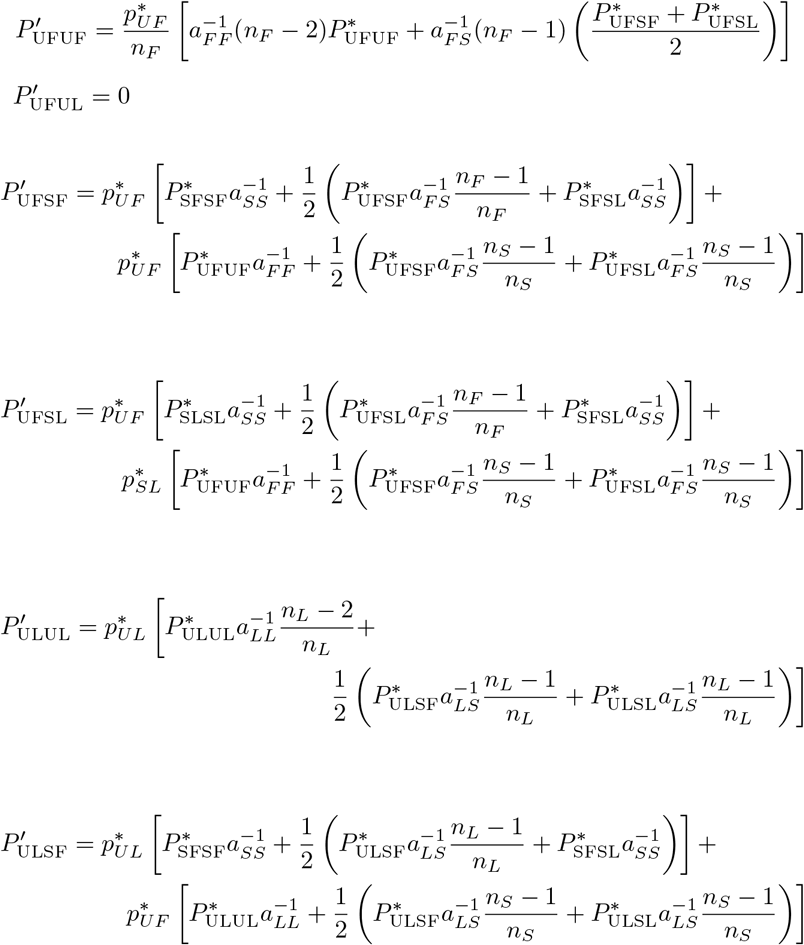

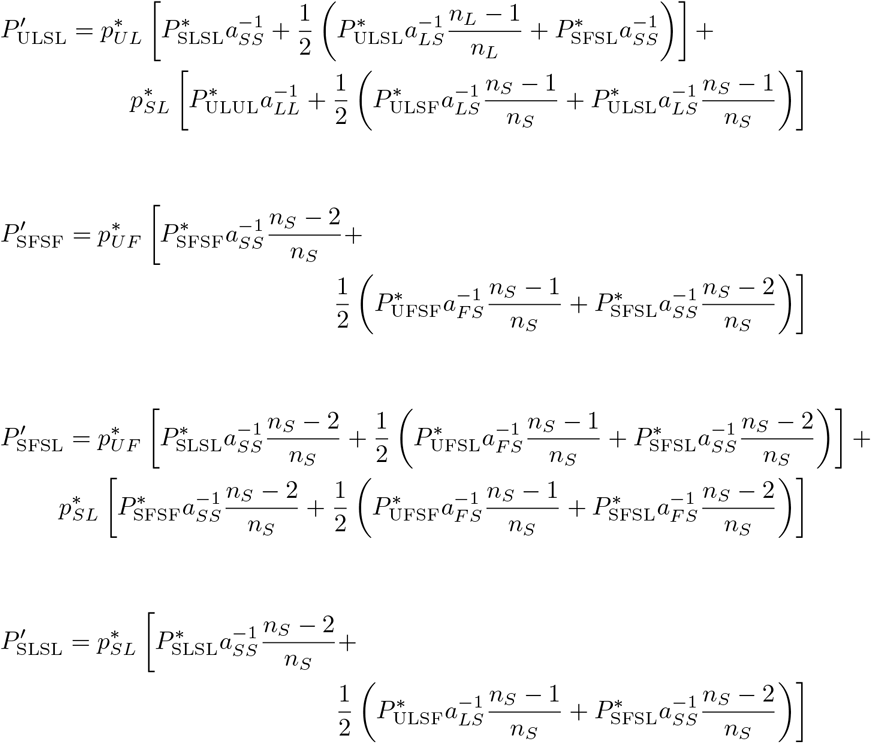

## 8.2 Self recognition

Assuming no pollen limitation, after migration and mating with self recognition, genotype frequencies are:

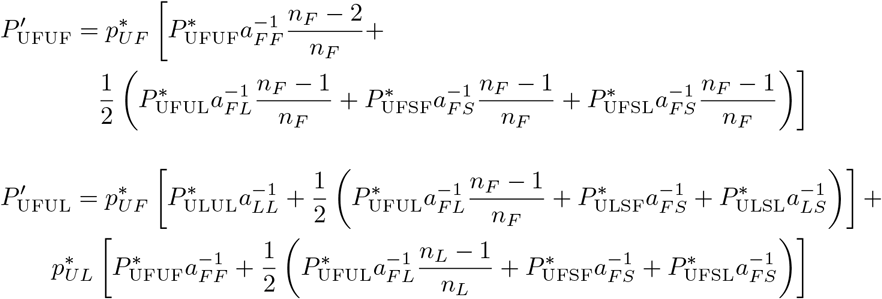

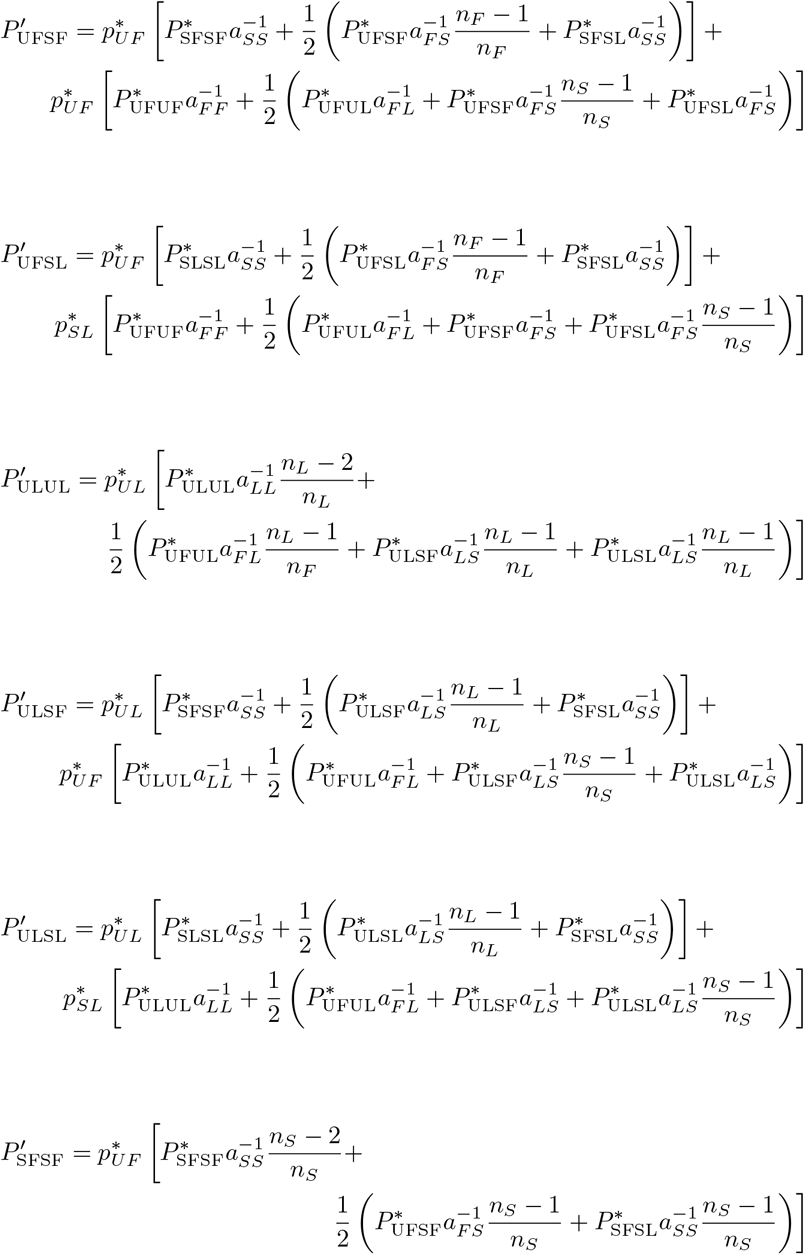

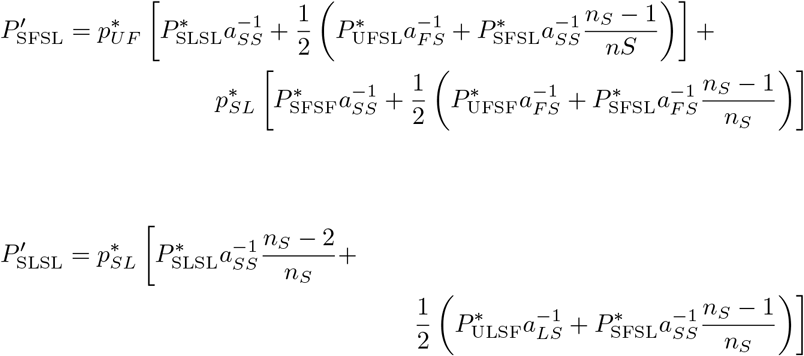

## References

K. Bod’ová, T. Priklopil, D. L. Field, N. H. Barton, and M. Pickup. Evolutionary pathways for the generation of new self-incompatibility haplotypes in a nonself-recognition system. Genetics, 209(3):861–883, 2018.

W. Boucher. A deterministic analysis of self-incompatibility alleles. Journal of Mathematical Biology, 31(2):149–155, 1993.

V. Castric, J. Bechsgaard, M. H. Schierup, and X. Vekemans. Repeated adaptive introgression at a gene under multiallelic balancing selection. PLoS Genetics, 4(8), 2008.

D. Charlesworth and B. Charlesworth. The evolution and breakdown of S-allele systems. Heredity, 43(1):41–55, 1979.

T. Dobzhansky. Studies on hybrid sterility. Zeitschrift für Zellforschung und mikroskopische Anatomie, 21(2):169–223, 1934.

V. E. Franklin-Tong, J. P. Ride, N. D. Read, A. J. Trewavas, and F. C. H. Franklin. The self-incompatibility response in *Papaver rhoeas* is mediated by cytosolic free calcium. The Plant Journal, 4(1):163–177, 1993.

S. Fujii, K.-i. Kubo, and S. Takayama. Non-self- and self-recognition models in plant self-incompatibility. Nature Plants, 2(9):1–9, 2016.

C. E. Gervais, V. Castric, A. Ressayre, and S. Billiard. Origin and diversification dynamics of self-incompatibility haplotypes. Genetics, 188(3):625–636, 2011.

A. Goldraij, K. Kondo, C. B. Lee, C. N. Hancock, M. Sivaguru, S. Vazquez-Santana, S. Kim, T. E. Phillips, F. Cruz-Garcia, and B. McClure. Compartmentalization of S-RNase and HT-B degradation in self-incompatible *Nicotiana*. Nature, 439(7078):805–810, 2006.

A. Harkness, E. E. Goldberg, and Y. Brandvain. Diversification or collapse of self-incompatibility haplotypes as outcomes of evolutionary rescue. BioRxiv, page 641613, 2019.

S. J. Hiscock. Pollen recognition during the self-incompatibility response in plants. Genome Bilogy, 3(2):reviews1004.1, 2002.

Z. Hua and T.-h. Kao. Identification and characterization of components of a putative *Petunia* S-locus F-box–containing E3 ligase complex involved in S-RNase–based self-incompatibility. The Plant Cell, 18(10):2531–2553, 2006.

S. Huang, H. S. Lee, B. Karunanandaa, and T.-h. Kao. Ribonuclease activity of *Petunia inflata* S proteins is essential for rejection of self-pollen. The Plant Cell, 6(7):1021–1028, 1994.

B. Igic and J. R. Kohn. Evolutionary relationships among self-incompatibility RNases. Proceedings of the National Academy of Sciences, 98(23):13167–13171, 2001.

K.-i. Kubo, T. Entani, A. Takara, N. Wang, A. M. Fields, Z. Hua, M. Toyoda, S.-i. Kawashima, T. Ando, A. Isogai, et al. Collaborative non-self recognition system in S-RNase–based self-incompatibility. Science, 330(6005):796–799, 2010.

K.-i. Kubo, T. Paape, M. Hatakeyama, T. Entani, A. Takara, K. Kajihara, M. Tsukahara, R. Shimizu-Inatsugi, K. K. Shimizu, and S. Takayama. Gene duplication and genetic exchange drive the evolution of S-RNase-based self-incompatibility in *Petunia*. Nature Plants, 1(1):14005, 2015.

Z. Lai, W. Ma, B. Han, L. Liang, Y. Zhang, G. Hong, and Y. Xue. An F-box gene linked to the self-incompatibility (S) locus of *Antirrhinum* is expressed specifically in pollen and tapetum. Plant molecular biology, 50(1):29–41, 2002.

D.-T. Luu, X. Qin, D. Morse, and M. Cappadocia. S-RNase uptake by compatible pollen tubes in gametophytic self-incompatibility. Nature, 407(6804): 649–651, 2000.

A. Muirhead. Consequences of population structure on genes under balancing selection. Evolution, 55(8):1532–1541, 2001.

H. Muller. Isolating mechanisms, evolution, and temperature. Biological Symposia, 6, 1942.

T. Nagylaki. The deterministic behavior of self-incompatibility alleles. Genetics, 79(3):545–550, 1975.

M. Pickup, Y. Brandvain, C. Fraïsse, S. Yakimowski, N. H. Barton, T. Dixit, C. Lexer, E. Cereghetti, and D. L. Field. Mating system variation in hybrid zones: facilitation, barriers and asymmetries to gene flow. New Phytologist, 224(3):1035–1047, 2019.

H. Qiao, H. Wang, L. Zhao, J. Zhou, J. Huang, Y. Zhang, and Y. Xue. The F-box protein AhSLF-S2 physically interacts with S-RNases that may be inhibited by the ubiquitin/26S proteasome pathway of protein degradation during compatible pollination in *Antirrhinum*. The Plant Cell, 16(3):582–595, 2004.

M. H. Schierup. The number of self-incompatibility alleles in a finite, subdivided population. Genetics, 149(2):1153–1162, 1998.

M. H. Schierup and X. Vekemans. Genomic consequences of selection on self-incompatibility genes. Current opinion in plant biology, 11(2):116–122, 2008.

M. H. Schierup, X. Vekemans, and F. B. Christiansen. Allelic genealogies in sporophytic self-incompatibility systems in plants. Genetics, 150(3):1187–1198, 1998.

M. R. Servedio and R. Bürger. The counterintuitive role of sexual selection in species maintenance and speciation. Proceedings of the National Academy of Sciences, 111(22):8113–8118, 2014.

W. Steiner and H.-R. Gregorius. Single-locus gametophytic incompatibility: the symmetric equilibrium is globally asymptotically stable. Journal of Mathematical Biology, 32(6):515–520, 1994.

P. Sun and T.-h. Kao. Self-incompatibility in *Petunia inflata*: The relationship between a self-incompatibility locus F-box protein and its non-self S-RNases. Plant Cell, 25(2):470–485, 2013.

M. K. Uyenoyama, Y. Zhang, and E. Newbigin. On the origin of self-incompatibility haplotypes: transition through self-compatible intermediates. Genetics, 157(4):1805–1817, 2001.

J. S. Williams, L. Wu, S. Li, P. Sun, and T.-H. Kao. Insight into S-RNase-based self-incompatibility in *Petunia*: recent findings and future directions. Frontiers in plant science, 6:41, 2015.

S. Wright. The distribution of self-sterility alleles in populations. Genetics, 24 (4):538, 1939.

